# Patch2MAP combines patch-clamp electrophysiology with super-resolution structural and protein imaging in identified single neurons without genetic modification

**DOI:** 10.1101/2023.03.20.533452

**Authors:** Dimitra Vardalaki, Trang L. D. Pham, Matthew P. Frosch, Garth Rees Cosgrove, Mark Richardson, Sydney S. Cash, Mark T. Harnett

## Abstract

Recent developments in super-resolution microscopy have revolutionized the study of cell biology. However, dense tissues require exogenous protein expression for single cell morphological contrast. In the nervous system, many cell types and species of interest – particularly human – are not amenable to genetic modification and/or exhibit intricate anatomical specializations which make cellular delineation challenging. Here, we present a method for full morphological labeling of individual neurons from any species or cell type for subsequent cell-resolved protein analysis without genetic modification. Our method, which combines patch-clamp electrophysiology with epitope-preserving magnified analysis of proteome (eMAP), further allows for correlation of physiological properties with subcellular protein expression. We applied Patch2MAP to individual spiny synapses in human cortical pyramidal neurons and demonstrated that electrophysiological AMPA-to-NMDA receptor ratios correspond tightly to respective protein expression levels. Patch2MAP thus permits combined subcellular functional, anatomical, and proteomic analyses of any cell, opening new avenues for direct molecular investigation of the human brain in health and disease.

## Introduction

Super-resolution microscopy enables the investigation of proteins with nanoscopic resolution^1^. Synthetic gel-based super-resolution imaging techniques, which include eMAP^2,3^ and ExM^4^, achieve ultrastructural imaging by coupling physical expansion of tissue with diffraction-limited microscopy. eMAP further provides the opportunity of highly multiplexed proteomic analysis, as processed tissue is amenable to multiple rounds of re-staining^2^. However, in dense, intermeshed tissue like the brain, exogenous protein expression (via genetically modified animals or viral transduction) is required to reconstruct cellular morphology and assign protein signals to correct subcellular compartments. This limits current super-resolution techniques to genetically-amenable cell types and species: for example, systematic analysis of protein distributions with cellular resolution is not currently possible in human brain tissue. This is a critical problem, as it is unclear which cells, which subcellular compartments (i.e. synapses, axons, dendrites), and which proteins produce malfunctions that mediate brain diseases like Alzheimer’s^5,6^, autism^7^, or epilepsy^8,9^, preventing development of effective therapies. We therefore developed Patch2MAP, a method that combines patch-clamp electrophysiology with eMAP for concomitant functional and super-resolution structural, and proteomic investigation of single cells in brain tissue from any species.

## Results

### Combining patch-clamp electrophysiology with epitope preserving magnified analysis of the proteome (eMAP)

To develop a system for morphological labeling of individual cells amenable to subsequent proteomic analysis, we capitalized on the well-known approach for full anatomical reconstruction via the biocytin-streptavidin complex^10^. This method consists of acquiring a whole-cell patch-clamp recording of a single cell, filling the patched neuron with biocytin, chemically fixing the brain slice, and utilizing the strong non-covalent bond between streptavidin and biotin to selectively introduce a fluorophore to the filled cell. This approach allows for saturated filling of neurons and morphological reconstruction of fine processes including dendritic spines and axons. To achieve super-resolution proteomic imaging of biocytin-filled cells, we modified the original eMAP protocol (Fig. 1a). We used cortical layer 5 pyramidal cells (L5 PCs) from acute brain slices of adult mice as our model. Initially, we were unable to recover the biocytin distribution and reconstruct the neuron after the tissue-gel hybridization step of eMAP, irrespective of when we introduced the streptavidin-Alexa Fluor 488 (before or after the hydrogel-tissue hybridization). Furthermore, we were unable to recover the morphology of the neuron even after using antibodies that target streptavidin. We reasoned that hydrogel-tissue hybridization may alter the biotin-streptavidin-fluorophore complex. We therefore added an additional step of fixation before the hydrogel-tissue hybridization. We then washed the tissue thoroughly to remove any excess formaldehyde and incubated it in a hydrogel monomer solution containing 30% acrylamide, 10% sodium acrylate, 0.1% bis-acrylamide, and 0.03% VA-044. This process allowed us to preserve the biocytin-streptavidin complex and visualize the morphology of the neuron in the tissue-gel hybrid at an intermediate expansion ratio (Fig. 1b). Using a vibratome, we re-sliced the original 300 μm thick slice (typical for slice physiology) into thinner slices (Fig. 1c) and incubated the resulting slices in a denaturation solution containing 6% SDS (w/v), 0.1M phosphate buffer, 50 mM sodium sulfite, and 0.02% sodium azide (w/v) in DI water. We then used commercially available antibodies to target synaptic proteins and subsequently expanded the tissue-gel in deionized (DI) water to visualize the ultrastructure of synapses (Fig. 1d, e).

**Fig. 1.**
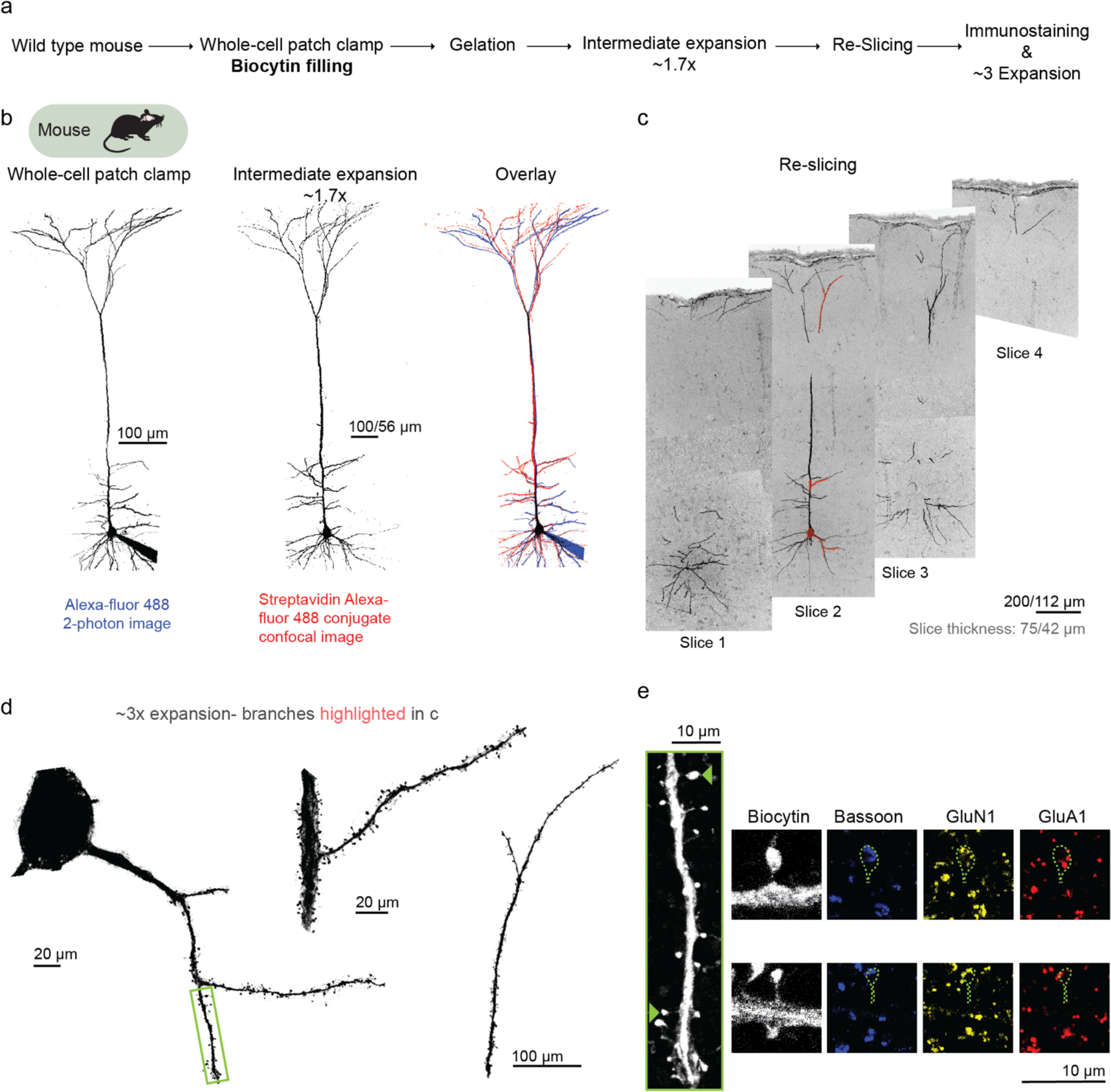
Patch2MAP enables super-resolution morphological and proteomic imaging in neural tissue without exogenous protein expression. **a**, Schematic of experimental tissue processing pipeline for full morphological labelling of individual neurons and subsequent cell-delineated super-resolution proteomic analysis. **b**, Left column: two-photon z-stack of a mouse L5 PC filled via somatic patch pipette with Alexa-488 and biocytin in an acute slice. Middle column: confocal z-stack of the same neuron at the stage of intermediate expansion. Strepatvidin Alexa-fluor 488 reveals biocytin. Right column: overlay of the two z-stacks. **c**, After intermediate expansion, the 300 μm-thick acute brain slice is resliced into 4 thinner slices. Confocal z-stacks show the 4 slices which contain the neuron from **b.** **d**, Confocal z-stacks of 3x-expanded soma and branches from the neuron in **b** and highlighted in **c.** **e**, Magnified view of basal branch from **d**. Green arrowheads indicate example spines shown to the right of the branch. From left to right: image of the cell-filling biocytin channel stained with Strepatvidin Alexa-fluor488, presynaptic protein Bassoon stained with anti-guinea pig-Alexa405 (blue), NMDAR subunit GluN1 stained with anti-mouse-Alexa555 (yellow), and AMPAR subunit GluA1 stained with anti-rabbit-Alexa647 (red).

### Patch2MAP enables super-resolution morphological reconstruction and multiplexed protein imaging of human neurons

To evaluate the applicability of Patch2MAP to human neurons, we acquired fresh human cortex from patients undergoing neurosurgical resection for treatment of epilepsy^11–14^. Blocks of cortical tissue were maintained in slicing aCSF, microdissected, and then sliced as was done for mouse cortex. We performed whole cell patch-clamp recordings of human cortical layer 2/3 pyramidal cells (L2/3 PCs) (Fig. 2a) and followed the same steps as in mice: fixation, gelation, and intermediate expansion followed by reslicing (Fig. 2b, c). Patch2MAP effectively produced bright and highly specific morphological and proteomic signals in human neurons and synapses (Fig. 2d, e, f, g). Furthermore, we showed that eMAP-processed human tissue gel can endure destaining and restaining of different proteins (Fig. 2g), demonstrating the suitability of this approach for the investigation of multiple proteins in each ultrastructural compartment. Thus, for the first time, Patch2MAP enables the concomitant ultrastructural investigation of human neuron morphology and protein distribution.

**Fig. 2.**
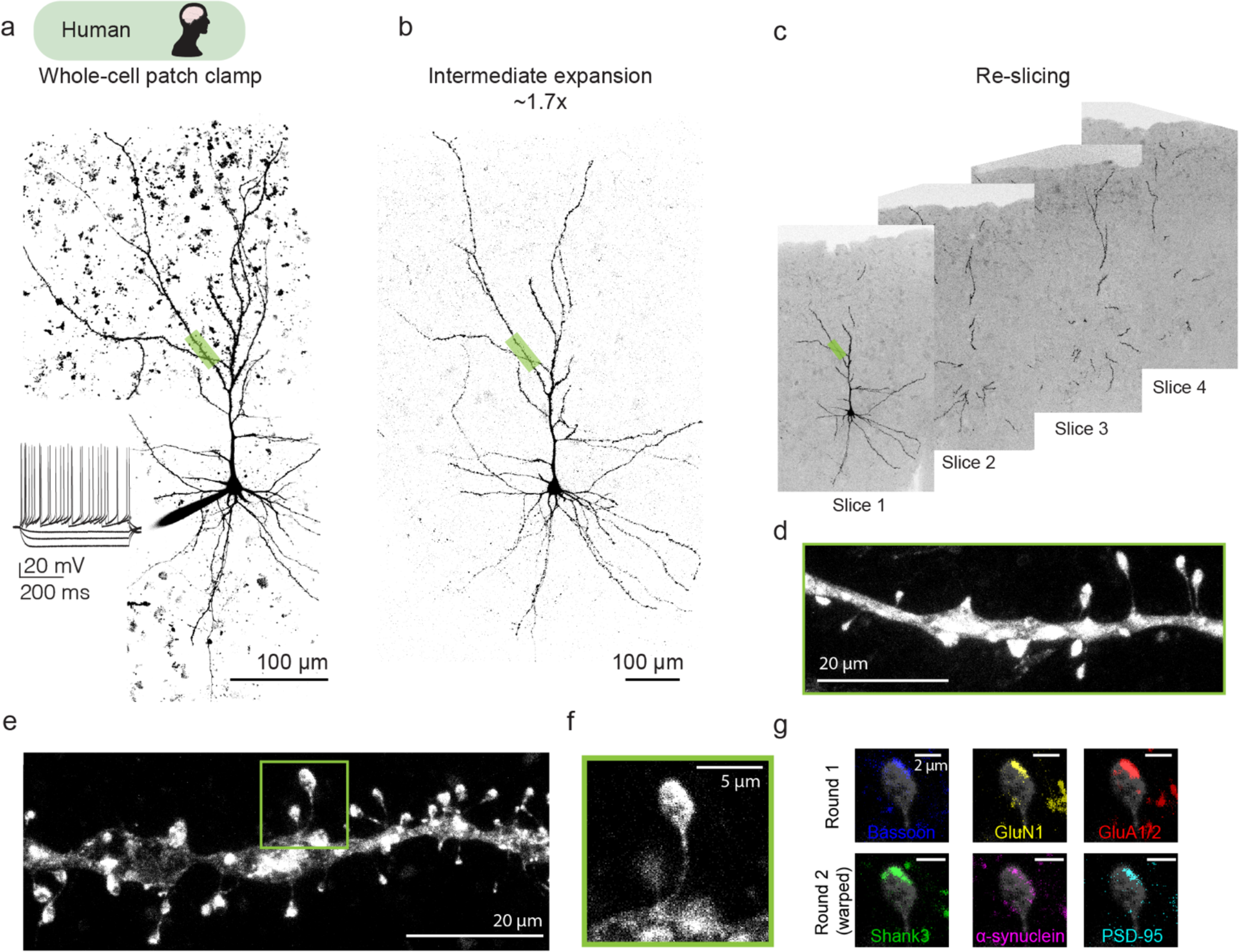
Patch2MAP enables super-resolution morphological and multiplexed proteomic imaging of human synapses. **a**, Two-photon z-stack and example somatic voltages in response to step current injections of a human L2/3 PC filled via somatic patch pipette with Alexa-488 and biocytin in an acute slice. **b**, Confocal z-stack of the same neuron at the intermediate expansion stage. Streptavidin Alexa-488 is used to show biocytin intracellular fill. **c**, Confocal images after re-slicing of the processed human brain slice into 4 thinner slices that contain the neuron shown in **b.** **d**, Confocal z-stacks of 3x expanded dendritic branch highlighted in **a, b, c.** **e**, Example basal branch from a different human neuron. Dendritic spine of interest indicated by green box. **f**, Magnified view of dendritic spine from **e.** **g**, Multi-round staining images of a Patch2MAP processed human tissue. Biocytin channels from both rounds were used as the reference for image registration. Round 1 (top): Bassoon (blue), GluN1 (yellow), and GluA1 & GluA2 (red). Round 2 (bottom): Shank3 (green), α-synuclein (magenta), and PSD-95 (cyan).

### Correlation of physiological properties with subcellular protein expression in human neurons

We took advantage of Patch2MAP to directly test how physiological properties correspond to subcellular protein expression. We targeted two classes of glutamate receptors, AMPA and NMDA receptors, at single spiny synapses in human cortical pyramidal neurons. AMPA and NMDA receptors play key roles in excitatory synaptic transmission and plasticity in the mammalian brain^15–18^. Despite more than 40 years of rigorous investigation, it is still unclear how the number of receptor proteins for a given synapse relates to functional synaptic strength.

We used whole cell patch-clamp recording and rapid two-photon glutamate uncaging to measure functional AMPA and NMDA receptor properties^19–23^ at individual spiny synapses in human neurons. The response of a given spine to glutamate uncaging relies on the uncaging laser power, as well as the uncaging location relative to the synapse of interest and the local delivery of the caged glutamate compound^24^. Additionally, spine neck resistance and dendritic filtering also affect the amplitude of synaptic responses recorded in the soma^25^. To account for these potential confounds, we measured the AMPA-to-NMDA receptor ratio (AMPA:NMDA) instead of absolute amplitude. AMPA:NMDA provides a measurement that is largely independent of uncaging parameters and dendritic filtering. We activated individual spines at a given dendritic branch with glutamate uncaging under control conditions in current-clamp mode to produce approximately physiologically-sized unitary uEPSPs (Fig. 3a). We then perfused the slice with DNQX and Mg^2+^-free aCSF and repeated the uncaging at the same spines (Fig. 3a, c, h). This allowed us to create a ratio of mostly AMPAR-mediated uEPSP amplitude to mostly NMDAR-mediated uEPSP amplitude. After the recording, we processed the tissue for subsequent super-resolution imaging, staining the tissue gel hybrid for GluA1 and GluA2 AMPA receptor subunits and the GluN1 NMDA receptor subunit. After ∼3x expansion of the tissue (Extended Data Fig. 1) we identified the dendritic branch containing the spines of interest and computed the AMPA:NMDA fluorescence signal intensity for every spine that was activated by glutamate uncaging (Fig. 3a, d, h, Extended Data Fig. 2). We found that the functional AMPA:NMDA correlates strongly and significantly with the protein AMPA:NMDA (Fig. 3e, i, j, Extended Data Fig. 2, 3, 4; n=76 spines, 11 cells, 4 human patients; Pearson correlation coefficient=0.65, P-value=1.6e-10). Our results validate that protein content can be reliably inferred by the signal intensity measured with Patch2MAP. Furthermore, protein content predicts synaptic transmission strength in human cortical neurons.

**Fig. 3.**
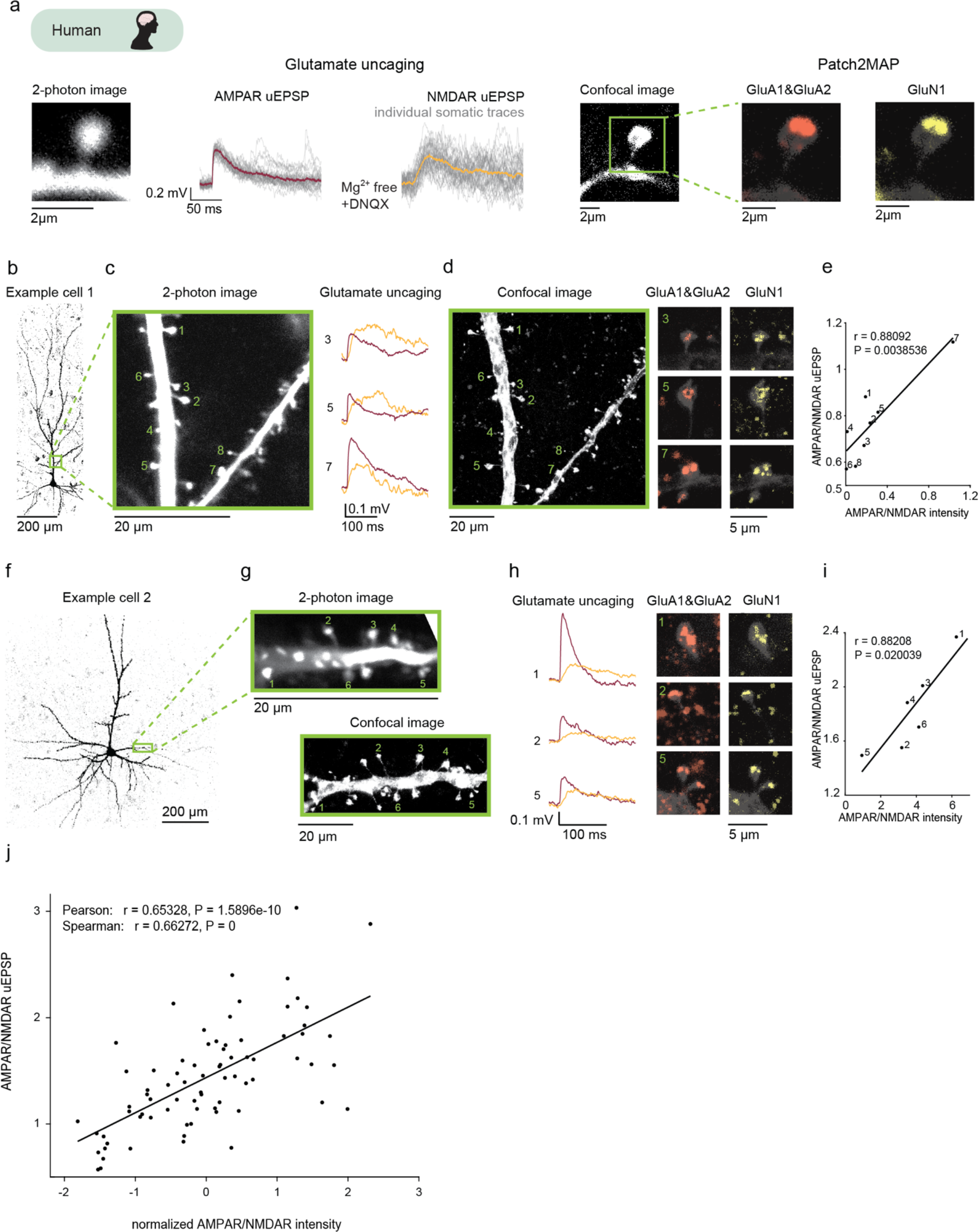
Protein content measured with Patch2MAP predicts synaptic transmission strength at human synapses. **a**, Right: two-photon image of a human spine filled via somatic patch pipette with Alexa-488 and biocytin in an acute slice. Middle: individual (grey) and average (colored) voltage traces recorded in current clamp mode at the soma in response to glutamate uncaging at the spine in aCSF (left) and after washing in Mg^2+^-free aCSF with 20 μM DNQX (right). Right: confocal images of the same spine after 3x expansion, from left to right: image of the cell-filling biocytin channel stained with Streptavidin Alexa-fluor 488, AMPAR subunit GluA1 and GluA2 stained with anti-rabbit-Alexa647 (red), and NMDAR subunit GluN1 stained with anti-mouse-Alexa555 (yellow). **b**, Confocal z-stack of a human PC at the intermediate expansion stage. Branch of interest is indicated by green box. **c**, Left: 2-photon image of the branch from **b** during whole cell patch-clamp. Numbers indicate the uncaging locations at individual spines. Right: example averaged voltage traces for spines 3,5 &7 in response to glutamate uncaging in control aCSF (red) and after washing in Mg^2+^-free aCSF with 20 μM DNQX (yellow). **d**, Right: confocal image of the same branch in **c** after ∼3x expansion through Patch2MAP processing. Left: AMPAR subunit GluR1 and GluR2 stained with anti-rabbit-Alexa647 (red) and NMDAR subunit NR1 stained with anti-mouse-Alexa555 (yellow) superimposed on cell-filling biocytin channel stained with Streptavidin Alexa-fluor 488 for spines 3,5 &7. **e**, Mean AMPAR/NMDAR uEPSP ratio versus AMPAR/NMDAR signal intensity ratio. Numbers correspond to the spines shown in **c, d** and **e**. The data are fit with a line of slope 0.4735; r: correlation coefficients, P: P value. **f**, As in **b** for another human PC. **g**, Top: 2-photon image of the branch from **f** in acute slice, Bottom: confocal image of the same branch after ∼3x expansion. **h**, As in **c** & **d** for example spines 1,2 & 3 from **g.** **i**, As in **e**. The data are fit with a line of slope 0.16576. **j**, Population correlation of mean AMPAR/NMDAR uEPSP ratio with normalized AMPAR/NMDAR signal intensity ratio (n=76 spines, 11 cells, 4 humans). The data are fit with a line of slope 0.33. (r) correlation coefficients, (P) P value.

## Discussion

We introduce a new method to perform super-resolution proteomic microscopy at identified subcellular structures without genetic modification by coupling patch-clamp electrophysiology with eMAP. Our approach provides a generally applicable framework to visualize subcellular morphology and protein content of neurons without exogenous protein expression. Moreover, we link biophysical properties to protein profiles at the level of individual synapses and further show that signal intensity of labeled proteins measured through Patch2MAP can be reliably used to infer the protein concentration. Our method enables the ultrastructural cell-delineated proteomic analysis of any neuron of any species, including humans, and paves the way for further studies in brain health and disease. While animal models are key for understanding human disease mechanisms and probing potential therapies^26^, there are significant challenges to creating useful models of human neurological disorders due the divergence of human brain organization and function^27–29^. Patch2MAP allows for the direct investigation of the human brain and the underlying mechanisms of neurological diseases.

## Acknowledgements

We thank Dae Hee Yun and Kwanghun Chung for sharing their expertise on eMAP. We thank Lukas Fisher, Marie-Sophie van der Goes, Jakob Voigts, and Courtney Yaeger for constructive criticism of the manuscript.

## Funding

Boehringer Ingelheim Fonds Foundation (D.V.)

NIH RO1NS106031 (M.T.H)

James W. and Patricia T. Poitras Fund at MIT (M.T.H.)

Klingenstein-Simons Fellow (M.T.H)

Vallee Foundation Scholar (M.T.H)

McKnight Scholar (M.T.H)

## Author Contributions

D.V. and M.T.H conceived of the project. D.V. designed experiments, performed electrophysiological recordings, processed the fixed tissue, performed super-resolution imaging, analyzed the data, and prepared figures. T.L.D.P. performed fixation, biocytin staining, gelation, denaturation, and immunostaining of the brain slices. M.R. and G.R.C. performed surgeries that resulted in human tissue. M.P.F. oversaw removal and parcellation of tissue as well as overall IRB aspects and regulatory aspects of the project with regard to human participants. S.S.C. helped in designing methods for acquiring human tissue and ensured that tissue was collected. M.T.H. supervised all aspects of the project and wrote the manuscript with D.V.

## Data and materials availability

All data are available upon request.

## Materials and Methods

### Animals

#### Human

Human (*Homo sapiens*) tissue was acquired through collaboration with the Massachusetts General Hospital (MGH) and the Brigham and Women’s Hospital (BWH). For MGH, tissue was obtained as ‘discarded tissue’ from neurosurgical patients in accordance with protocols approved by the Massachusetts General Hospital Internal Review Board (IRB). Patients consent to surgery and a subset of the resected tissue was considered discarded tissue. Under MGH’s IRB-approved protocol, such discarded tissue was available for this specific research project for use without explicit patient consent. For BWH, the protocol was approved by the IRB, and patients provided consent prior to the surgery. At both institutions, non-essential samples were extracted by the supervising neuropathologist per protocol.

Patients were women or men aged 23–71 years. Additional patient information is included in Extended Data Table 1. Samples were not allocated to distinct experimental groups and information about the patient was not available until after data acquisition and analysis.

#### Mouse

All animal procedures were done in compliance with the NIH and Massachusetts Institute of Technology Committee on Animal care guidelines. We used C57BL/6 mice (Charles River Laboratories). Male and female mice were used in approximately equal numbers for all experiments at 8-10 weeks of age. Mice were kept on a 12-hour light/dark cycle and had unrestricted access to food and water.

### Brain slice preparation

#### Human slice preparation

Resected human tissue was considered discarded tissue after being examined by neuropathologists whose main objective was to ensure there was adequate tissue for diagnostic purposes. Neocortical tissue was obtained from the lateral anterior temporal and frontal lobe in patients undergoing resection for medically-intractable epilepsy. The neocortical tissue displayed no known abnormalities at the level of MRI scans, gross inspection, and subsequent microscopic examination as part of the standard neuropathologic assessment of the tissue. Patients undergoing resective surgery were primarily maintained under general anesthesia with propofol and remifentanil or sufentanil. Some cases utilized inhaled anesthetics, such as isoflurane or sevoflurane. For induction of general anesthesia, paralytic agents including rocuronium or succinylcholine as well as fentanyl were typically used. Resection usually occurred within 90 minutes of the start of the procedure.

After resection, tissue was placed in ice-cold cutting solution containing (in mM): sucrose 165, sodium bicarbonate 25, potassium chloride 2.5, sodium phosphate 1.25, calcium chloride 0.5, magnesium chloride 7, glucose 20, HEPES 20, sodium pyruvate 3, and sodium ascorbate 5, 295-305 mOsm, equilibrated with 95% O2 and 5% CO_2_. Samples were transported in sealed conditions for ∼20 minutes before being transferred to freshly oxygenated solution. Pia and surface blood vessels that would obstruct slicing were removed. Slicing was performed with a Leica VT1200S vibratome in ice-cold cutting solution. 300 μm-thick slices were incubated for ≥ 30 minutes at 36°C in recovery solution containing (in mM): sodium chloride 90, sodium bicarbonate 25, potassium chloride 2.5, sodium phosphate 1.25, calcium chloride 1, magnesium chloride 4, glucose 20, HEPES 20, sodium pyruvate 3, and sodium ascorbate 5, 295-305 mOsm, equilibrated with 95% O_2_ and 5% CO_2_. Slices were then stored at room temperature until use. Incubation solutions were replaced every ∼8 hours and recordings were performed up to 34 hours after slicing.

#### Mouse slice preparation

Coronal brain slices (300 μm) containing the primary visual cortex (V1) were prepared from 8- to 10-week-old C57BL/6 mice. Animals were deeply anesthetized with isoflurane prior to decapitation. The brain was removed and sliced with a vibratome (Leica) in ice-cold slicing solution containing (in mM): sucrose 90, NaCl 60, NaHCO_3_ 26.5, KCl 2.75, NaH_2_PO_4_ 1.25, CaCl_2_ 1.1, MgCl_2_ 5, glucose 9, sodium pyruvate 3, and ascorbic acid 1, saturated with 95% O_2_ and 5% CO_2_. Slices were incubated in artificial cerebrospinal fluid (aCSF) containing (in mM): NaCl 120, KCl 3, NaHCO_3_ 25, NaH_2_PO_4_ 1.25, CaCl_2_ 1.2, MgCl_2_ 1.2, glucose 11, sodium pyruvate 3, and sodium ascorbate 1, saturated with 95% O_2_ and 5% CO_2_ at 35.5 °C for 25-30 min and then stored at room temperature.

Recording aCSF containing (in mM): sodium chloride 120, potassium chloride 3, sodium bicarbonate 25, sodium phosphate monobasic monohydrate 1.25, calcium chloride 1.2, magnesium chloride 1.2, glucose 11, sodium pyruvate 3, and sodium ascorbate 1, 302-305 mOsm, saturated with 95% O_2_ and 5% CO_2_.

### Patch2MAP protocol

#### Patch-clamp recording and Biocytin filling

Patch-clamp recordings were performed from the soma of pyramidal neurons at 34–36 °C in recording aCSF containing (in mM): sodium chloride 120, potassium chloride 3, sodium bicarbonate 25, sodium phosphate monobasic monohydrate 1.25, calcium chloride 1.2, magnesium chloride 1.2, glucose 11, sodium pyruvate 3 and sodium ascorbate 1, 302–305 mOsm, saturated with 95% O_2_ and 5% CO_2_.

An Olympus BX-61 microscope with infrared Dodt optics and a 60x water immersion lens (Olympus) was used to visualize cells. Whole-cell current-clamp recordings were performed in bridge mode with a Dagan BVC-700 amplifier with bridge fully balanced. Current and voltage signals were filtered at 10 kHz and digitized at 20 kHz. Patch pipettes were prepared with thin-wall glass (1.5 O.D., 1.1 I.D.). Pipettes had resistances ranging from 3 to 7 MΩ and the capacitance was fully neutralized prior to break in. Series resistances ranged from 7-17 MΩ. The intracellular solution contained (in mM): potassium gluconate 134, KCl 6, HEPES buffer 10, NaCl 4, Mg_2_ATP 4, NaGTP 0.3, phosphocreatine di(tris) 14, 0.1 Alexa 488 (Invitrogen), 5.2 Biocytin (Invitrogen).

During the patch-clamp recording biocytin diffused from the patch pipette to the neuron. A recording time of at least 10 minutes was sufficient to completely fill the axon and the dendrites of large pyramidal neurons in mouse neocortical layers 5 and human neocortical layers 2 and 3. Longer times may be needed for larger neurons or for studies performed at room temperature. Upon completion of the electrophysiological recording, outside-out patch configuration was established by slowly retracing the recording pipette in voltage clamp mode under visual control. The capacitance and input resistance were monitored and loss of capacitive transients with the collapse of the current responses to a straight line ensured resealing of the cell membrane.

The tissue was then processed through our modified protocol for eMAP^2^, which allowed recovery of biocytin signals for subsequent morphological reconstruction.

#### Fixation and Biocytin Labelling

300 μm thick slices were transferred in 15 ml Falcon tubes containing 10 ml of 4% PFA in phosphate-buffered saline (PBS) at 4°C and incubated overnight. Slices were incubated in 50 ml Falcon tubes containing 40 ml of PBS for at least 24h at 4°C. Slices were transferred subsequently into 48-well plates containing 200 μl of washing solution (PBS containing 0.1% (w/v) Triton X-100(PBST) and 0.02% (w/v) sodium azide) on 37°C shaker for 4h. PBST was at least once exchanged before adding 20 μl of Streptavidin, Alexa Fluor™ 488 Conjugate (S32354, Thermo Fisher, 0.2% (w/v)) and slices were incubated on 37°C shaker overnight. Slices were washed 4 times for 10 min each in PBST. This washing process was repeated twice with an interval of 2 hours and slices were transferred to 4% PFA in phosphate-buffered saline (PBS) at 4°C and incubated for 8-12h. Slices were switched to PBS at 4°C for at least 36h before performing confocal imaging with excitation at 488 nm to image biocytin filled-neurons.

#### Gelation

Slices were incubated in eMAP hydrogel monomer solution (30% acrylamide (A9099, MilliporeSigma, St. Louis, MO, USA), 10% sodium acrylate (408220, MilliporeSigma), 0.1% bis-acrylamide (161-0142, Bio-Rad Laboratories, Hercules, CA, USA), and 0.03% VA-044 (w/v) (Wako Chemicals, Richmond, VA, USA) in PBS) at 4°C overnight. Then, slices were carefully placed between two glass slides using Blu-Tack adhesive (Bostik, Essendon Fields, Victoria, Australia) and with the help of ∼290 μm thick homemade spacers. The empty space was filled with additional hydrogel monomer solution. The glass slide setup was then placed inside a 50 ml Falcon tube, which was subsequently placed inside Easy-Gel (LifeCanvas Technologies, Cambridge) with nitrogen gas at 37°C for 3 hours. After gelation, the cartridge was disassembled, and excess gel was trimmed from around the lateral edges of the sample using a razor blade. The resulting tissue-gel hybrid sample was put in PBS with 0.02% sodium azide at 37°C shaker overnight for hydration.

#### Re-slicing

The tissue gel hybrid was first expanded by 1.7x. Confocal imaging was performed to identify the neuron and to determine slicing orientation. A thin layer of Krazy Glue (Elmer’s products, inc. 460 Polaris Parkway Westerville, OH 4308) was applied on a 10 mm non-orienting specimen disc (14048143399, Leica Biosystems, Germany). An 300 μm thick slice was placed (neuron closer to the slicing surface) on the glue in the specimen disc and light pressure is applied with a cover slip to make sure uniform application of glue. The tissue gel hybrid was then resliced using a vibratome (VT1200, Leica Biosystems, Germany) in 75 μm (44 μm original) thick slices. From the resulting slices, the ones that contain the neuron of interest are further trimmed with a razor blade to final dimensions of ∼3mm*3mm*75μm.

#### Denaturation & Immunostaining

The slice was incubated in a denaturation solution [6% SDS (w/v), 0.1 M phosphate buffer, 50 mM sodium sulfite, and 0.02% sodium azide (w/v) in DI water] at 37°C shaker for 6h. Subsequently, the samples were transferred in washing solution (PBS containing 0.1% (w/v) Triton X-100 (PBST) and 0.02% (w/v) sodium azide) at 37°C shaker overnight. The washing solution was exchanged at least once before adding the primary antibodies. The slice was incubated with primary antibodies (typical dilution 1:20) in PBST at 37°C shaker overnight, followed by washing in PBST at 37°C for 6 hours with 4 solution exchanges. The same process was repeated for secondary antibodies (typical dilution 1:10)

#### Mounting & image acquisition

Before each imaging session the slice was incubated in DI water for 1m to reach final expansion (3x). Then the expanded specimen was placed between a petri dish and a glass-bottom Willco dish (HBSB-5030; WillCo Wells, Amsterdam, The Netherlands) using glass coverslips as spacers. To prevent the samples from drying up during image acquisition, the void space around the samples was filled with DI water. Subsequently, the samples were imaged using a Zeiss LSM 900-AS microscope system using a 63x 1.2 NA water immersion objective or a Leica TCS SP8 microscope system using a 63× 1.2 NA water immersion objective.

#### Multi-round staining

Previously stained tissues were incubated in denaturation solution for 6 hours at 37°C, followed by 10 m at preheated 95°C denaturation solution to remove bound antibodies. Tissue specimens were then washed in PBST at 37°C overnight. PBST was exchanged at least twice. The samples were imaged on the microscope to confirm the complete loss in the signal from the antibodies. Tissue was then immunostained as described above.

#### Expansion factor measurement

To measure the expansion factor, we imaged the same cells at 3 time points: first, during 2-photon imaging of patched cell, then at intermediate expansion of the gel-tissue hybrid, and finally after final expansion (13 human slices). For each cell we calculated the mean expansion ratio for each cell from at least 3 measurements among preserved neuronal structures along the x,y dimensions.

### Glutamate uncaging

A two-photon laser scanning system (Prairie Technologies Ultima) with dual galvanometers and two ultrafast pulsed lasers beams (Mai Tai DeepSee lasers) were used to simultaneously image and uncage glutamate. One path was used to image Alexa 488 at 920 nm. The other path was used to photolyse MNI-caged L-glutamate (Tocris) at 720 nm. Stock MNI-glutamate solutions (50 mM) were freshly diluted in Mg^2+^ -free aCSF to 10 mM and a Picospritzer (General Valve) was used to focally apply the MNI-caged L-glutamate via pressure ejection through a large glass pipette above the slice. Laser beam intensity was independently controlled with electro-optical modulators (model 350-50; Conoptics). Emitted light was collected by GaAsP photomultipliers. Uncaging dwell time was 0.2 ms. A passive 8X pulse splitter in the uncaging path was used to reduce photodamage^30,31^. Experiments were not further analyzed if diffuse signs of photodamage were detected (increase in basal fluorescence, loss of transient signals, and/or persistent depolarization).

#### Measurement of AMPA-to-NMDA receptor ratio at individual spines

Uncaging locations were positioned in close vicinity of spines (<0.5 μm) from the tip of individual spine heads in the radial direction. 6-12 spines were individually stimulated at each branch. The uncaging stimulus was delivered in each spine separately (using a stimulus interval of 500 ms). Unitary uEPSPs at each spine were evoked 20 to 40 times and responses were averaged. Mg^2+^ free aCSF containing 20 μM DNXQ was then washed on for at least 15 minutes. The same uncaging protocol was repeated at the same spines. Care was taken to maintain the initial uncaging locations and focal plane throughout the experiment.

### Measurement of uEPSP amplitudes

The AMPAR-mediated and NMDAR-mediated uEPSP amplitude for each spine was measured offline using custom-written MATLAB code. Briefly, the peak amplitude of each recorded uEPSP was measured during the peak window (50 ms post glutamate uncaging). The width of the window was chosen with consideration of both AMPAR and NMDAR dynamics. Baseline potential was calculated as the average voltage in the 50 ms preceding the evoked uEPSP and was then subtracted from the measurement of the peak uEPSP to provide the individual uEPSP amplitude. Individual uEPSP amplitudes (20-40) were averaged to provide AMPAR-mediated and NMDAR-mediated uEPSPs amplitude for each spine.

### Image analysis

#### Identification of spines of interest

The identification of spines of interest is based on: 1) gross morphology (identification of the branch) and 2) spine imaging (identification of the spines). Both are achieved by comparing the 2-photon (physiology step) and confocal (expansion step) images in terms of geometry, branching points, and relative location to other branches/spines. Examples for identifying the same branches before and after expansion are shown in Fig. 1&2, and for identifying the same spines before and after the expansion are shown in Fig. 3c, d, g. The spines, which do not have an unambiguous match in the expanded tissue (Extended Data Fig. 3b, c) are excluded (see also **Correlation analysis**).

#### Measurement of signal intensity in AMPAR and NMDAR channels

Dendritic spines were analyzed in the biocytin channel using Fiji software. We used a custom-written macro code that first applies a median blur (2 pixels) in the biocytin image and then converts the tip of individual spines to binary masks by thresholding the resulting biocytin image. The rest of the analysis was performed using custom-written MATLAB code. Each image had 4 channels (bassoon, biocytin, NMDAR, AMPAR) and multiple z-slices (z-step of 0.66 μm). The signal intensity in each of the four channels was calculated at every z-slice as the intensity difference of the mask containing the structure of interest and the background. The background intensity was calculated by using the same mask but at random x-y locations of the image to account for the effect of size in the measured intensity among dendritic protrusions. The z-slices that contained the spine of interest were defined as the z-slices where the biocytin intensity signal for a given spine was greater than the median biocytin signal intensity for the same spine from every z-slice +1 SD. The signal intensity for the AMPAR and NMDAR channels for a given spine was calculated as the sum of the signal intensity in all the z-slices that contained the respective spine, while the controls (Extended Data Fig. 2) were calculated by 1000 random combinations of the same number of slices that did not contain the respective spine.

#### Z-score normalization of signal intensity

In order to pool the data of all cells in a single plot (Fig. 3j, Extended Data Fig. 3e), we z-scored the AMPAR:NMDAR signal intensity among each subset of cells, which were treated with the same antibodies. The polyclonal antibodies used in our experiments (GluA1 subunit,182 003 Sysy and GluA2 subunit,182 103 Sysy) are a complex mixture of several antibodies and commercial vials differ from each other. Normalization allows to account for differences of the inherent signal intensity of secondary antibodies.

## Correlation analysis

We analyzed 11 dendritic branches (91 spines and 11 cells from 4 humans) where we performed the uncaging experiment and then reconstructed the full length of the dendritic segments with Patch2MAP. From the 2-photon data, 4 spines (4/91) were excluded because they showed signs of photodamage. From the eMAP data, 5 spines (5/91) were excluded because the super-resolution morphological imaging revealed that 2 spines instead of 1 were targeted by the 2-photon glutamate uncaging, and 6 spines (6/91) were excluded because the uncaging location did not correspond to a distinct spine in the expanded branch (Extended Data Fig. 3). The signal intensity for those was calculated either in the photodamaged spine or in one of the two spines or in the adjacent dendritic branch (shaft synapses). These exclusions did not change the results presented in the main figures (Extended Data Fig. 3e).

## References

1. Sahl, S. J., Hell, S. W. & Jakobs, S. Fluorescence nanoscopy in cell biology. Nature Reviews Molecular Cell Biology vol. 18 685–701 (2017).

2. Park, J. et al. Epitope-preserving magnified analysis of proteome (eMAP). Sci. Adv. 7, 1–14 (2021).

3. Ku, T. et al. Multiplexed and scalable super-resolution imaging of three-dimensional protein localization in size-adjustable tissues. Nat. Biotechnol. 34, 973–981 (2016).

4. Chen, F., Tillberg, P. W. & Boyden, E. S. Optical imaging. Expansion microscopy. Science 347, (2015).

5. Wang, J., Gu, B. J., Masters, C. L. & Wang, Y. J. A systemic view of Alzheimer disease - Insights from amyloid-β metabolism beyond the brain. Nat. Rev. Neurol. 13, 612–623 (2017).

6. Götz, J., Bodea, L. G. & Goedert, M. Rodent models for Alzheimer disease. Nat. Rev. Neurosci. 19, 583–598 (2018).

7. Lord, C., Elsabbagh, M., Baird, G. & Veenstra-Vanderweele, J. Autism spectrum disorder. Lancet 392, 508–520 (2018).

8. Barker-Haliski, M. & Steve White, H. Glutamatergic mechanisms associated with seizures and epilepsy. Cold Spring Harb. Perspect. Med. 5, 1–15 (2015).

9. Finnema, S. J. et al. Imaging synaptic density in the living human brain. Sci. Transl. Med. 8, 1–10 (2016).

10. Liu, F., Zhang, J. Z. H. & Mei, Y. The origin of the cooperativity in the streptavidin-biotin system: A computational investigation through molecular dynamics simulations. Sci. Rep. 6, (2016).

11. Testa-Silva, G. et al. High Bandwidth Synaptic Communication and Frequency Tracking in Human Neocortex. PLoS Biol. 12, (2014).

12. Berg, J. et al. Human neocortical expansion involves glutamatergic neuron diversification. Nature 598, 151–158 (2021).

13. Beaulieu-Laroche, L. et al. Allometric rules for mammalian cortical layer 5 neuron biophysics. Nature 600, 274–278 (2021).

14. Beaulieu-Laroche, L. et al. Enhanced Dendritic Compartmentalization in Human Cortical Neurons. Cell 175, 643-651.e14 (2018).

15. Henley, J. M. & Wilkinson, K. A. Synaptic AMPA receptor composition in development, plasticity and disease. Nat. Rev. Neurosci. 17, 337–350 (2016).

16. Meldrum, B. S. Glutamate as a neurotransmitter in the brain: Review of physiology and pathology. J. Nutr. 130, 1007–1015 (2000).

17. Malenka, R. C. & Nicoll, R. A. NMDA-receptor-dependent synaptic plasticity: multiple forms and mechanisms. Trends Neurosci. 16, 521–527 (1993).

18. Huganir, R. L. & Song, I. Regulation of AMPA receptors during synaptic plasticity. TRENDS Neurosci. 25, 578–588 (2002).

19. Lafourcade, M. et al. Differential dendritic integration of long-range inputs in association cortex via subcellular changes in synaptic AMPA-to-NMDA receptor ratio. Neuron 1–15 (2022) doi:10.1016/j.neuron.2022.01.025.

20. Yaeger, C. E., Vardalaki, D., Brown, N. J. & Harnett, M. T. Dendritic compartmentalization of input-specific integration and plasticity rules across cortical development. bioRxiv 2022.02.02.478840 (2022).

21. Malinow, R. & Malenka, R. C. AMPA receptor trafficking and synaptic plasticity. Annu. Rev. Neurosci. 25, 103–126 (2002).

22. Magee, J. C. & Cook, E. P. Somatic EPSP amplitude is independent of synapse location in hippocampal pyramidal neurons. Nat. Neurosci. 3, 895–903 (2000).

23. Bekkers, J. M. & Stevens, C. F. NMDA and non-NMDA receptors are co-localized at individual excitatory synapses in cultured rat hippocampus. Nature 341, 230–233 (1989).

24. Ellis-Davies, G. C. R. Two-Photon Uncaging of Glutamate. Front. Synaptic Neurosci. 10, 1–13 (2019).

25. Rall, W. Theoretical significance of dendritic trees for neuronal input-output relations. Neural Theory Model. ed. R.F. Reiss. Stanford Univ. Press. (1964).

26. van der Worp, H. B. et al. Can Animal Models of Disease Reliably Inform Human Studies? PLOS Med. 7, e1000245 (2010).

27. Nestler, E. J. & Hyman, S. E. Animal models of neuropsychiatric disorders. Nat. Neurosci. 13, 1161–1169 (2010).

28. Rice, J. Animal models: Not close enough. Nature 484, S9–S9 (2012).

29. Hartung, T. Thoughts on limitations of animal models. Park. Relat. Disord. 14, 83–85 (2008).

30. Ji, N., Magee, J. C. & Betzig, E. High-speed, low-photodamage nonlinear imaging using passive pulse splitters. Nat. Methods 5, 197–202 (2008).

31. Harnett, M. T., Xu, N. L., Magee, J. C. & Williams, S. R. Potassium channels control the interaction between active dendritic integration compartments in layer 5 cortical pyramidal neurons. Neuron 79, 516–529 (2013).

