## Supplementary material for "Patch2MAP combines patch-clamp electrophysiology with super-resolution structural and protein imaging in identified single neurons without genetic modification": Supplemetal Fig1-4, Supplemental Table 1

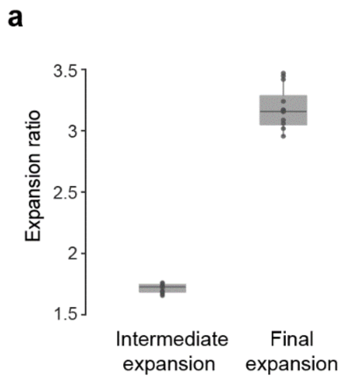


**Extended Data Fig. 1: Expansion ratio**
Expansion ratios for human cells at the intermediate expansion step and at the final expansion step (n=13 cells, 5 humans)

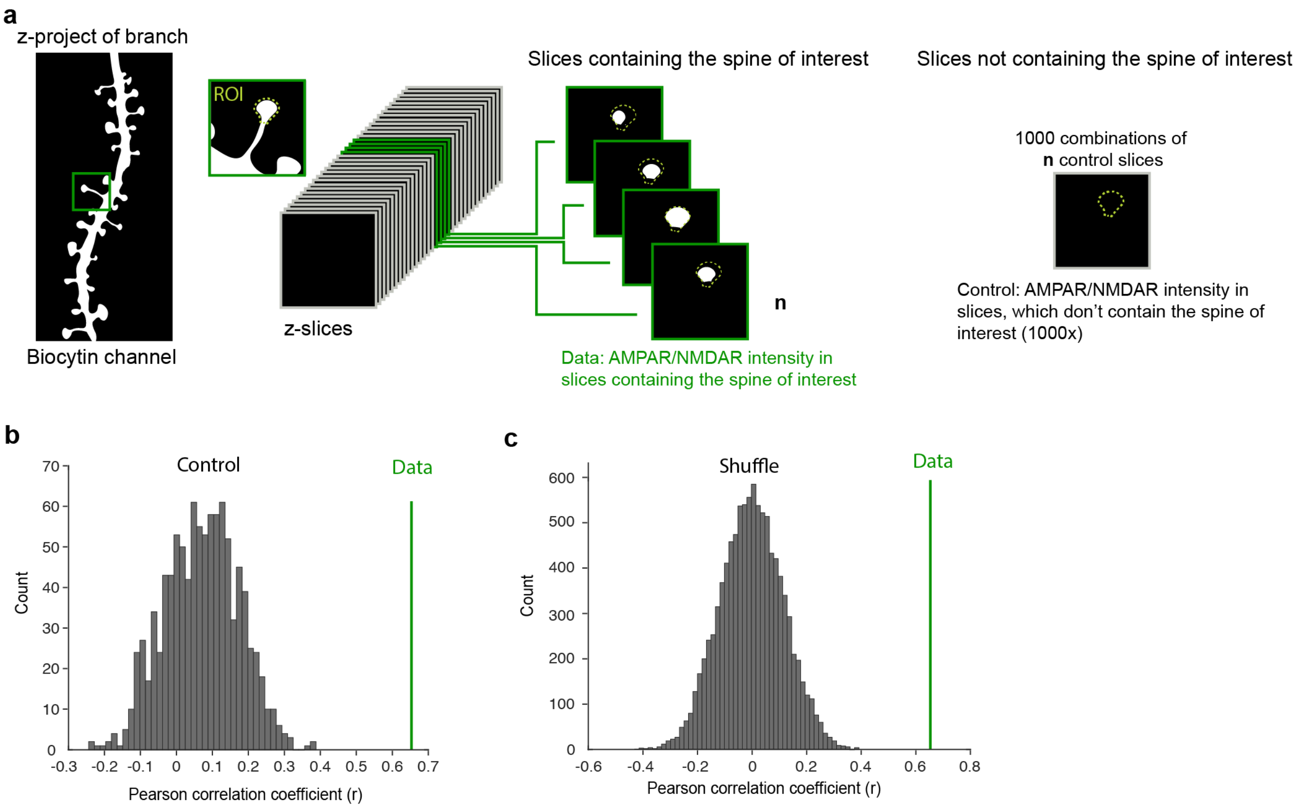


**Extended Data Fig. 2. Description of analysis and control experiments**
**a,** Schematic of analysis pipeline for calculating the AMPAR/NMDAR intensity for each synapse and the corresponding control.
**b,** Correlation values of electrophysiological to antibody intensity ratio from data are above the 100$^{th}$ percentile (z-score= 5.6622) of 1000 control measurements of AMPA/NMDA intensity.
**c,** Same as **b** for 10000 randomly drawn pairs of electrophysiological AMPA/NMDA ratios and antibody AMPA/NMDA signal intensities from experimental data (z-score= 5.6403).

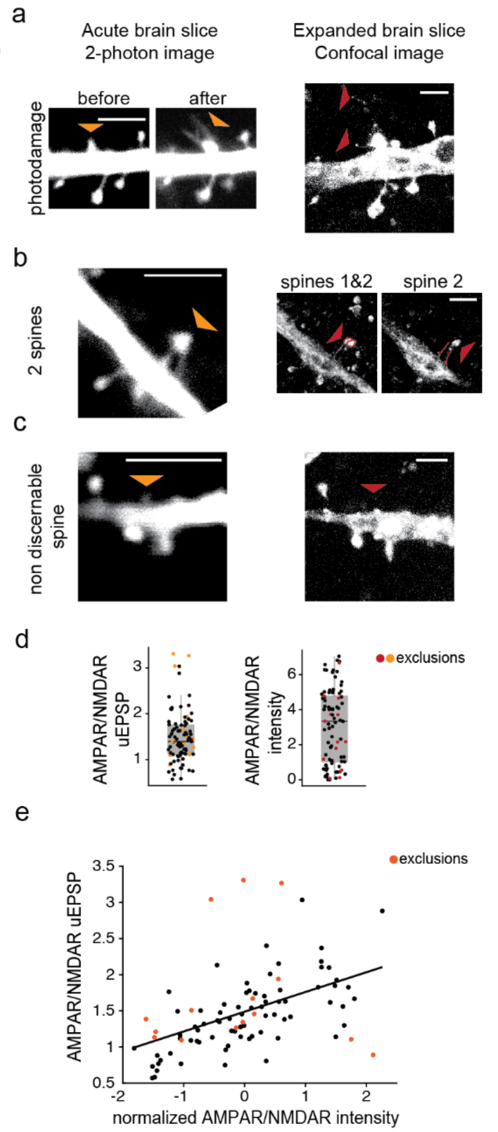


**Extended Data Fig. 3: Examples of exclusion criteria**
**a,** Example of a spine with signs of photodamage. Left: 2-photon images of the spine (orange arrowhead) before and after the initiation of uncaging experiment. Right: confocal image of the same spine in the expanded tissue. Note the spinules arising from the spine (red arrowheads).
**b,** Example of a spine where super-resolution revealed that two spines where targeted by glutamate uncaging instead of one. Left: 2-photon image of the targeted spines (orange arrowhead); Right: confocal images of the same spines in the expanded tissue showing the distinct necks (red lines) and heads (red arrowheads) of the two spines.
**c,** Example of uncaging location that did not correspond in a distinct spine in the expanded branch. Left: 2-photon image of the target location (orange arrowhead). Right: confocal image of the same branch in the expanded tissue. No corresponding spine is found (red arrowhead)
**d,** Distribution of all the AMPAR/NMDAR for uEPSPs (left) and antibody signal intensity (right). Exclusions highlighted in color. Box plot represents median and IQR with whiskers extending to the most extreme points not considered outliers.
**e,** Dots, correlation of mean AMPAR/NMDAR uEPSP ratio and normalized AMPAR/NMDAR signal intensity ratio (see Fig. 3j).

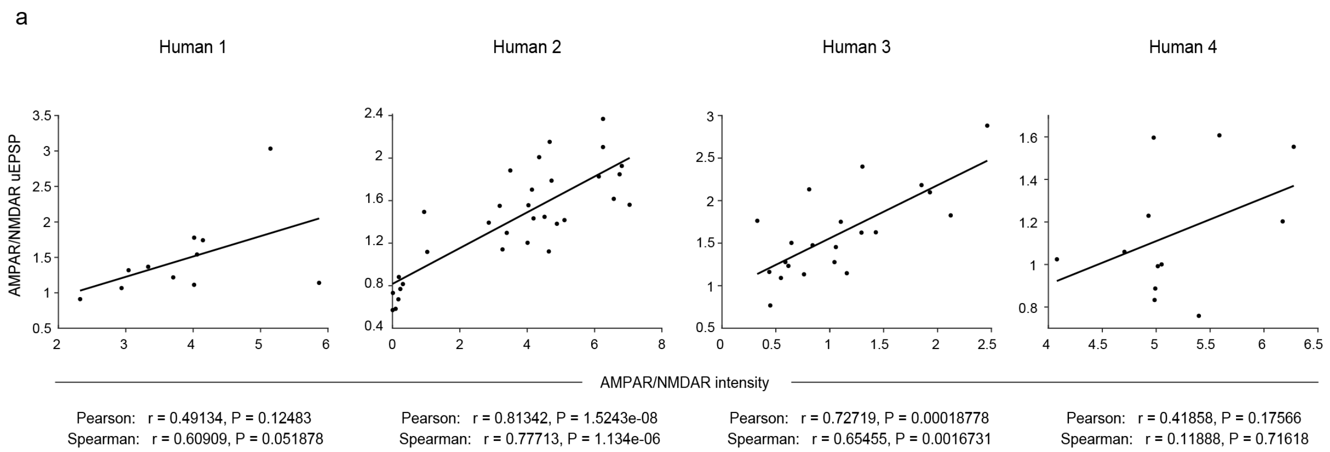


**Extended Data Fig. 4: Correlation of AMPAR/NMDAR uEPSP and AMPAR/NMDAR signal intensity per human subject.**

Correlation of mean AMPAR/NMDAR uEPSP ratio and AMPAR/NMDAR signal intensity ratio. r: correlation coefficients, P: P value.

| Sex | Age | Handness | Age at epilepsy onset | Brain region | Antiepileptic drugs (pre-surgery) |
|---|---|---|---|---|---|
| M | 71 | R | 60 | R TL | BRV, LTG |
| M | 33 | R | 31 | R FL | BRV, LCM, LOR |
| F | 23 | L | 6 | R ATL | LTG, LOR |
| M | 24 | R | 6 | R ATL | LCM, OCB |
| F | 37 | R | 16 | R TL | LCM, OCB |

**Extended Data Table 1: Patient information**

Male (M), female (F), right (R), left (L), temporal lobe (TL), frontal lobe (FL), anterior temporal lobe (ATL), brivaracetam (BRV), lamotrigine (LTG), lacosamide (LCM), lorazepam (LOR), oxcarbazepine (OCB)
